# HiNT: a computational method for detecting copy number variations and translocations from Hi-C data

**DOI:** 10.1101/657080

**Authors:** Su Wang, Soohyun Lee, Chong Chu, Dhawal Jain, Geoff Nelson, Jennifer M. Walsh, Burak H. Alver, Peter J. Park

## Abstract

The three-dimensional conformation of a genome can be profiled using Hi-C, a technique that combines chromatin conformation capture with high-throughput sequencing. However, structural variations (SV) often yield features that can be mistaken for chromosomal interactions. Here, we describe a computational method HiNT (**Hi**-C for copy **N**umber variation and **T**ranslocation detection), which detects copy number variations and inter-chromosomal translocations within Hi-C data with breakpoints at single base-pair resolution. We demonstrate that HiNT outperforms existing methods on both simulated and real data. We also show that Hi-C can supplement whole-genome sequencing in SV detection by locating breakpoints in repetitive regions.

## BACKGROUND

The Hi-C assay provides genome-wide identification of chromatin interactions, thereby enabling systematic investigation of the three-dimensional genome architecture and its role in gene regulation [1]. Hi-C data have been used, for example, to characterize topologically associated domains (TADs), which are megabase-sized local chromatin interaction domains within which genomic loci interact with higher frequency [2–4]. Characterization of genome organization using Hi-C data has enhanced our understanding of a number of biological processes, such as X-inactivation [2, 5], cell cycle dynamics [6], and tumor progression [7].

However, it has been shown that structural variations (SVs) can confound the interpretation of Hi-C data [6, 8–11]. For example, when there is copy number increase, the observed number of sequencing reads that correspond to chromosomal interactions in that region will be larger than expected, not because there is greater frequency of interaction but because there are multiple copies of that region. Similarly, when there is an inter-chromosomal translocation, the reads that correspond to interactions between the translocated segment and its proximal regions will be inflated, but this should not be mistaken for changes in interaction frequency.

One approach to mitigate the impact of SVs on the Hi-C interaction map is to first identify SVs using whole-genome sequencing (WGS) data and then use that information to adjust the Hi-C map. Although a great deal of progress has been made in WGS-based SV detection [12, 13], the use of WGS data requires additional sequencing and analysis expertise. Furthermore, SV breakpoints within repetitive regions, which are often genomic SV hotspots, cannot be easily detected from WGS due to low mappability [14]. Indeed, Hi-C and WGS data are complementary in SV detection: as Hi-C read pairs span genomic distances from base pairs to megabases, they enable detection of breakpoints in repetitive regions when one read of a read pair maps to a repetitive region and the other maps to a surrounding mappable region (Supp. Fig. 1).

Here, we present HiNT (**Hi**-C for copy **N**umber variation and **T**ranslocation detection), an algorithm for detection of copy number variations (CNVs) and inter-chromosomal translocations in Hi-C data. Based on simulated data and comparisons to variants identified in WGS, HiNT outperforms existing computational methods both in sensitivity and false discovery rate (FDR). HiNT also provides translocation breakpoints at single base-pair resolution, a feature not available in existing methods that utilize only Hi-C data. Furthermore, HiNT supports parallelization, utilizes efficient storage formats for interaction matrices, and accepts multiple input formats including raw FASTQ, BAM, and contact matrix. HiNT is available at https://github.com/parklab/HiNT.

## RESULTS

### Overview of HiNT

HiNT has three main components. HiNT-PRE performs preprocessing of Hi-C data and computes the contact matrix, which stores contact frequencies between any two genomic loci. HiNT-CNV and HiNT-TL start with a Hi-C contact matrix and predict copy number segments and inter-chromosomal translocations, respectively (Supp. Fig. 2).

HiNT-PRE aligns read pairs to the genome using BWA-MEM [15] and creates a Hi-C contact matrix. The matrix is constructed from normal read pairs (non-chimeric reads that map uniquely to the genome) as well as *unambiguous* chimeras [16] (Fig. 1A). The latter is a product of Hi-C ligation and is defined as a read pair in which one chimeric read is split into locus A and locus B and the other read is uniquely mapped to locus B (Fig. 1A). All other read pairs containing split reads are defined as *ambiguous* chimeras [16], which will be used for translocation breakpoint detection (Fig. 1A).

**Figure 1.**
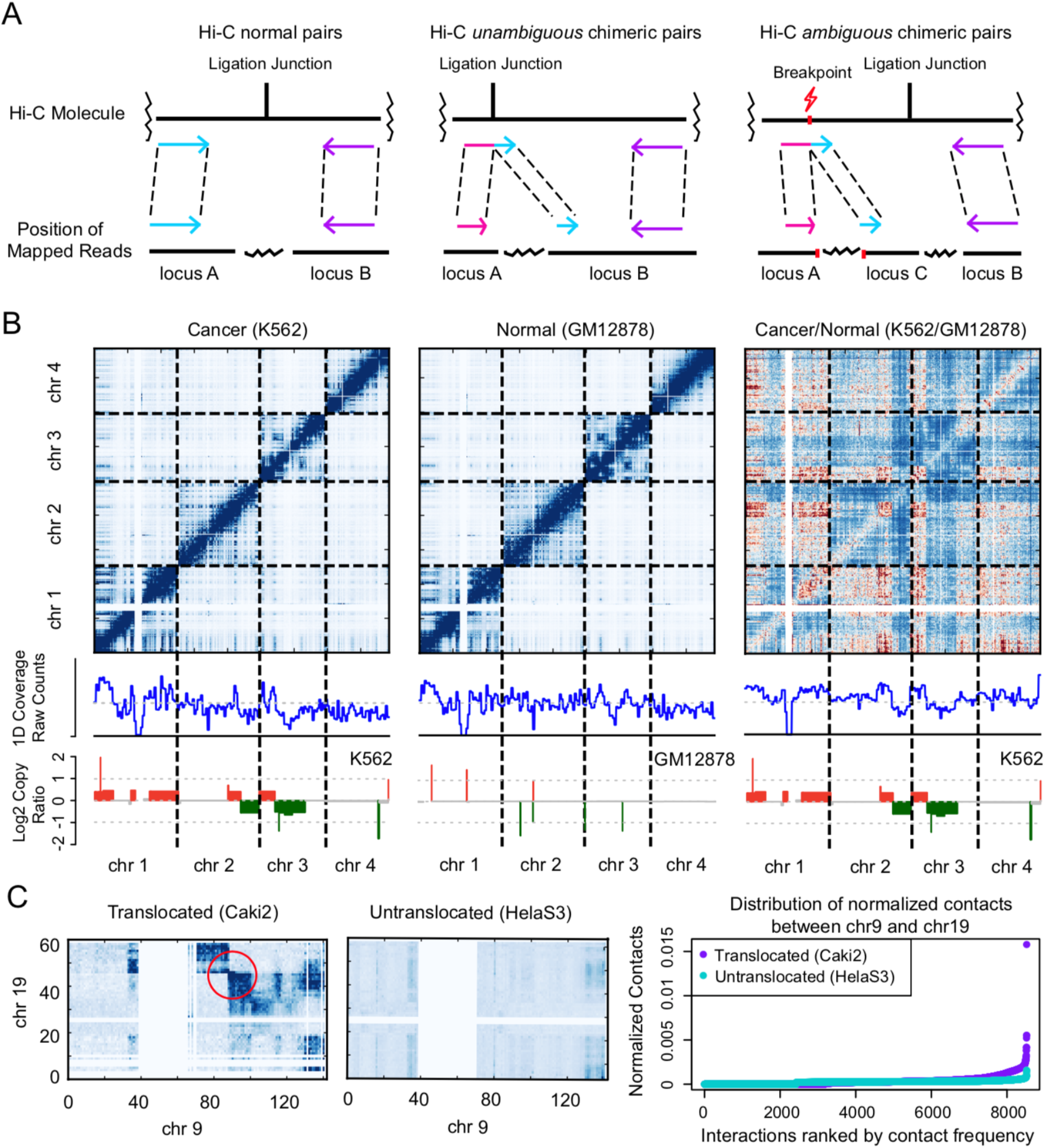
Illustration of HiNT. **A**, Hi-C read pairs are classified into normal pairs (left panel), *unambiguous* chimeric pairs (middle panel), and *ambiguous* chimeric pairs (right panel). Hi-C *unambiguous* chimeric pairs are the product of Hi-C ligations in which one read crosses the ligation junction and thus maps to both locus A and locus B, while the other normal read maps only to locus B. Hi-C *ambiguous* chimeric pairs are often caused by structural variations, with one read maps to both locus A and locus C, while the other read maps to locus B. **B**, Copy number information is reflected in the Hi-C 1D coverage profile after Hi-C biases are removed by normalizing the K562 Hi-C contact matrix with the GM12878 Hi-C contact matrix. The copy number profile (log2 ratios) estimated from WGS data is shown in the bottom row for comparison. **C**, Comparison of the Hi-C contact matrix between chr9 and chr19 in samples with and without translocations. The distribution of normalized contact frequencies are higher in the sample with translocation (purple dots) than in the sample without (cyan dots). Contact frequencies were calculated in 1Mb bins in chr9 and chr19.

HiNT-CNV (Supp. Fig. 2) first creates a one-dimensional (1D) coverage profile across the genome by calculating row or column sums of the contact matrix at a fixed resolution, e.g., 50kb. These sums should be correlated with the copy number across the bins since they correspond to the strength of interaction of that region with all other regions. It is critical to use the *unnormalized* contact matrix here because the matrix-balancing normalization (setting the sum of each row or column to be 1), which is the most widely used Hi-C normalization approach, removes not only biases but also copy number information. The next step is to perform further adjustment to remove other biases that are inherent in the Hi-C experiments, such as GC content, mappability, restriction site frequency, etc. In Fig. 1B, we see that, without additional adjustment, the 1D profiles for K562 (human chronic myelogenous leukemia cell line; known to have high genomic instability) and GM12878 (human lymphoblastoid cell line) show similarity to each other but not with the copy number profiles estimated from WGS. However, when we remove Hi-C internal biases in K562 by using GM12878 as a control (Fig. 1B, right), the 1D coverage profile becomes highly correlated with the (ploidy-adjusted) copy ratios estimated from WGS data. This result shows that proper normalization is essential in extracting copy number information from Hi-C data. Given that an appropriate control is often unavailable, HiNT-CNV uses a generalized additive model to remove the biggest sources of bias: GC content, mappability, and restriction fragment length [17]. The boundaries of CNV segments are determined using the BIC-seq segmentation algorithm, which utilizes the Bayesian information criterion to identify regions with enriched or depleted read counts [18].

HiNT-TL (Supp. Fig. 2) detects translocations by analyzing normalized inter-chromosomal interaction matrices. In general, contact probabilities between two regions on the same chromosome decrease monotonically with distance, and inter-chromosomal interactions are considerably less frequent compared to intra-chromosomal ones. When an inter-chromosomal translocation occurs, we expect the contact probabilities in two opposite quadrants around the breakpoint to be elevated to the levels observed for adjacent chromosomal regions (Fig. 1C). Thus, HiNT-TL identifies candidate translocated chromosomal pairs based on the presence of high contact probabilities and their unequal distribution. To identify exact breakpoints, HiNT-TL first identifies the breakpoint regions with a coarse 100kb resolution from the 1D profiles (see Methods). HiNT-TL then uses Hi-C *ambiguous* chimeric reads located within these regions to refine breakpoints to single base-pair resolution.

### CNVs predicted by HiNT from Hi-C are consistent with those identified from WGS

To predict CNVs, we first calculate the coverage profile throughout the genome at 50kb resolution. We then correct for Hi-C biases such as GC content, mappability, and the number of restriction sites (given a fixed bin size, the number of expected fragments depends on the number of cut sites by the restriction enzyme used). To model the non-linear correlation between 1D coverage and biases observed (Supp. Fig. 3), we use a generalized additive model (GAM) with the Poisson link function. GAM is an ideal framework here, as it allows non-parametric fitting with relaxed assumptions on the relationship between predictor and response variables. The copy number information is extracted from regression residuals by the following equation: *log (Coverage’) = s_1_(GCcontent)* + *s_2_(Mappability) + s_3_(NumberOfRestr.Sites) + ε* where *s_i_* (*i*=1,2,3)(•) is an unspecified function estimated for each predictor variable and *ε* is the regression residual. The model fits better for GM12878 (*R*^2^ = 0.798) than for K562 (*R*^2^ = 0. *631*), since K562 is known to have more SVs.

To evaluate CNVs identified from Hi-C, we compare the log2 copy ratios along the genome from the model above with those estimated from WGS. For K562, we see that copy number alterations are prevalent and that the log ratios from Hi-C and WGS are mostly concordant (Fig. 2A, Supp. Fig. 4A; Spearman correlation = 0.82). For GM12878, the correlation is lower (Spearman correlation = 0.21) because there are very few CNVs in this cell line, and the existing small ones are detected only from WGS (Supp. Fig. 4B, Supp. Fig. 5A). The copy ratios fluctuate more in the Hi-C profile relative to WGS data (Fig. 2A, Supp. Fig. 5A) due to the different read depth and possibly due to Hi-C biases that may not have been captured by our model. When the copy number log ratios are segmented using BIC-seq [18], the concordance between the platforms is striking (top two rows in Fig. 2B), with ~85% and 92% of the large (>2Mb) segments from Hi-C overlapping those from WGS in K562 and GM12878 cells, respectively (Fig. 2C, Supp. Fig. 5D; our definition of overlap is described in Supp. Fig. 5C). Collectively, our analysis suggests that HiNT is a reliable tool for identifying large-scale CNVs in both cancer and normal Hi-C data.

**Figure 2.**
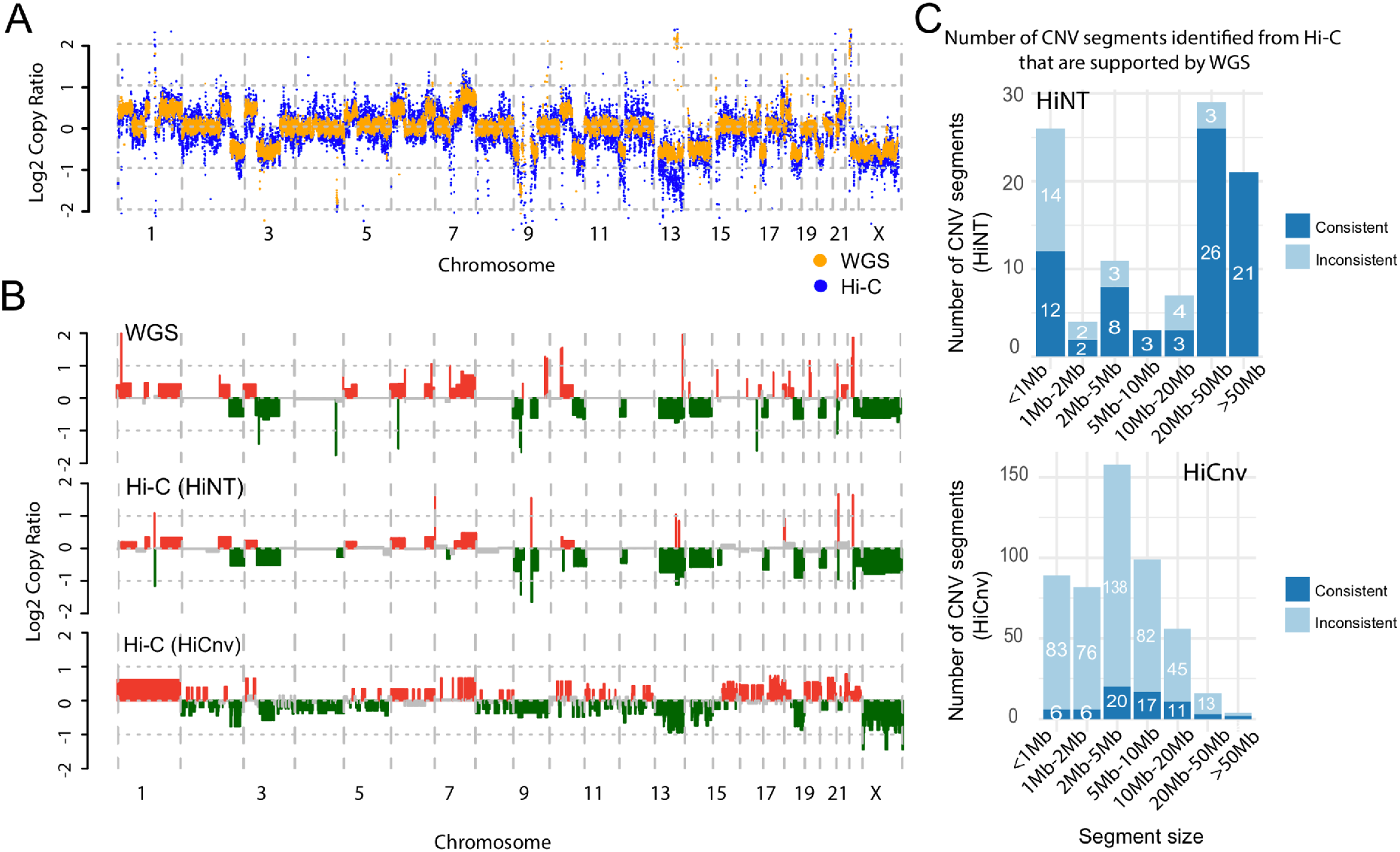
Copy number inference in K562 cells by HiNT. **A**, Comparison of log2 copy ratios calculated using regression residuals from Hi-C (blue) and using read coverage from WGS (orange). **B**, Comparison of CNV profiles from Hi-C and WGS after segmentation. Red, green and grey bars represent copy gain, copy loss, and copy neutral regions, respectively. **C**, The number of CNV segments detected from Hi-C by HiNT (upper) and HiCnv (lower) that are also supported by WGS. The overlap criteria for consistency are shown in Supp. Fig. 5C.

### HiNT outperforms HiCnv for identifying CNVs from Hi-C data

HiCnv is a computational tool developed to infer copy number from normalized Hi-C coverage [19]. It employs smoothing by kernel density estimation followed by a Hidden Markov Model; however, it requires a baseline chromosome copy number from WGS or karyotyping to determine the true copy number of each chromosome. To evaluate the performance of HiCnv, we examine the concordance of CNVs identified from HiCnv to those detected from WGS. Surprisingly, the copy number log ratios along the genome are largely discordant, with a Spearman correlation of 0.67 in K562 and 0.1 in GM12878 (Supp. Fig. 4C-D). Moreover, only ~15% of the large segments detected by HiCnv overlap those identified from WGS in K562 (Fig. 2C); the overlap is even smaller for GM12878 cells (Supp. Fig. 5D).

In addition, input to HiCnv must be either HiC-Pro [20] output or a SAM file, which is then converted to HiC-Pro format, incurring high computational cost for terabyte-scale datasets. For example, 3 billion read pairs result in a ~600GB BAM file, and the required SAM format is at least 4-fold larger than BAM format in size. In contrast, HiNT-PRE accepts FASTQ and BAM files, and generates the Hi-C contact matrix in hic [16, 21] or cool [22] format, which serves as the input to HiNT-CNV. Both hic and cool are efficient and widely-used formats for genomic interaction matrices. Taken together, HiNT-CNV outperforms this existing tool in detecting CNVs in both cancer and normal cells in both accuracy and usability.

### HiNT accurately identifies translocated chromosomal pairs

Translocations modify the 3D organization of the genome, and they will be incorrectly identified as long-range interactions in Hi-C data if they are not accounted for properly. To first study their impact on Hi-C interaction maps, we developed a simulation scheme to recapitulate the effect of translocations, encompassing homozygous/heterozygous and balanced/unbalanced translocations. A balanced translocation is an even exchange of segments between chromosomes without genetic information gain or loss; an unbalanced translocation involves a loss or gain of chromosome segments. As observed in previous studies [19, 23, 24], a balanced translocation forms a ‘butterfly’ appearance in the chromosomal interaction map (Fig. 3A and Fig. 3B middle, marked by red circles). In contrast, an unbalanced translocation only has a single block (Fig. 3A and Fig. 3B, right column, marked by red circles) [23]. Detection of intra-chromosomal translocations are complicated by the presence of chromatin structures such as TADs and loops. Therefore, we focus on identification of inter-chromosomal translocations.

**Figure 3.**
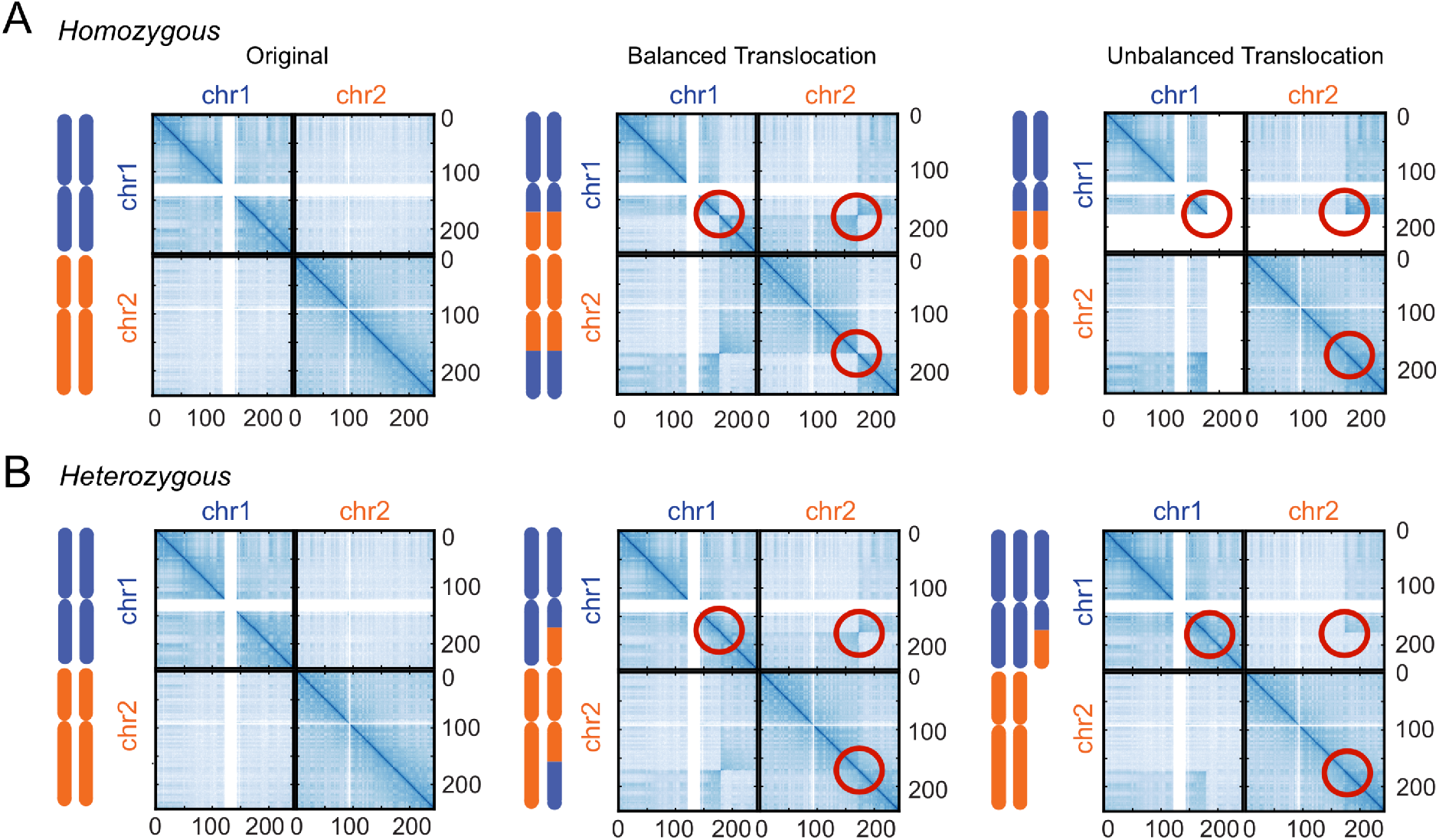
Simulation translocations in Hi-C data. **A**, Homozygous cases. **B**, Heterozygous cases. An example of a translocation involving two chromosomes is illustrated. The three columns correspond to original matrix, with balanced translocation, and unbalanced translocation, respectively. Circles highlight the features introduced by the translocations.

Our method is based on detection of two characteristics. First, the contact frequencies should be distributed unevenly around the translocation breakpoint. For this, we utilize the Gini index, a statistical measure of distribution initially used to quantify income inequality in economics [25]. To compute this index, we estimate the cumulative distribution of contact frequencies in each square of the interaction map (we use 1Mb x 1Mb) and determine how much it deviates from a linear increase. A high index corresponds to more uneven distribution of interaction strength. Second, the maximum interaction level surrounding the breakpoint should be high for a translocation. Regions without a translocation but with a high noise level may satisfy the first criterion of uneven contact frequencies, but their maximum interaction level would not be large. Combining the two features (interaction level and evenness), we define the rank product score as *RP_i_* = (*R_gini,i_/n*)*S* * (*R_mif,i_/n*), where *R_gini,i_* and *R_mif,i_* are the ranks of matrix *í* based on Gini index and maximum interaction frequency, respectively, and *n* is the total number of inter-chromosomal interaction matrices.

The rank product score performs well in simulated data, separating the translocated and non-translocated cases in nearly all cases (Supp. Fig. 6). For real data, we found that direct application of the rank product was insufficient, due to the various factors that are not captured by the normalization step, e.g., the A/B compartment effect and the increased interactions between small chromosomes or between sub-telomeric regions. To eliminate such biases, we created a background interaction matrix by averaging the matrices from five normal cell lines (Supp. Table 1, see Methods) and used it to normalize the original matrix. In Fig. 4A, we show three examples of chromosomal pairs in K562 data whose scores change as a result of the additional normalization. In the first case (chr1-chr21), the score does not change significantly; in the second case (chr1-chr18), the score increases so that a translocation is now called; and in the third case (chr16 – chr19), the score decreases so that a mistaken call is avoided. Using the chromosomal pairs reported in the literature or validated by FISH experiments [4, 24] as true positives, we see that the adjusted matrix results in an increased prediction accuracy, as measured by the area under the curve (AUC) (Fig. 4B). As visualized in Fig. 4C, the previously observed biases are effectively reduced by the normalization, allowing for better delineation of translocations (Supp. Fig. 6, Supp. Fig. 7A-C).

**Figure 4.**
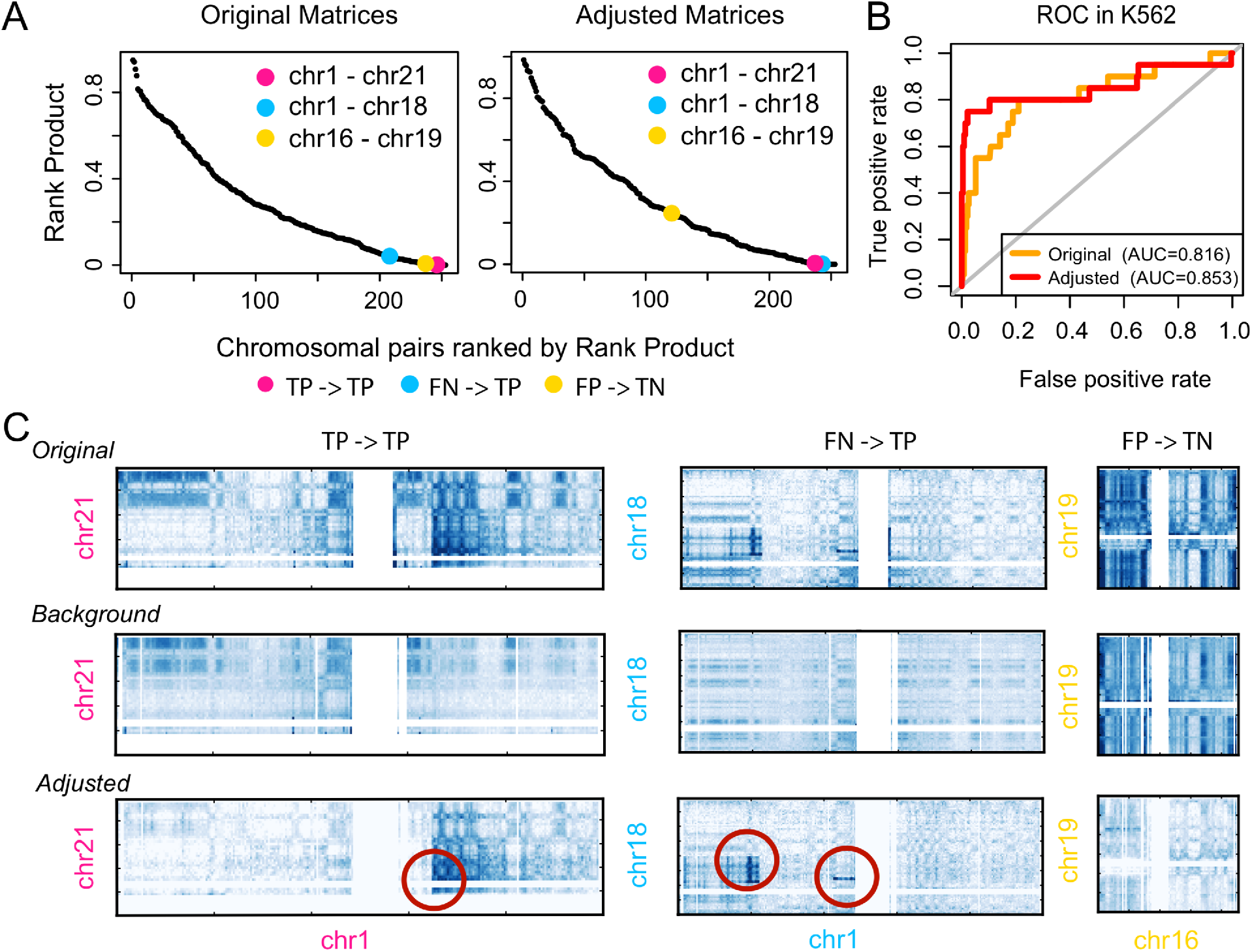
Accurately identification of translocated chromosomal pairs by HiNT. **A**, The distribution of the rank product scores for all chromosomal pairs in K562 before (left) and after (right) adjustment by background subtraction. Chromosomal pairs in pink and blue correspond to two FISH-validated translocation pairs (chr1, chr21) and (chr1, chr18); the one in yellow corresponds to a chromosome pair (chr16, chr19) without translocation. After matrix adjustment, the blue pair now has a lower score and the yellow pair has a higher score, as desired. TP: True Positive, TN: True Negative, FN: False Negative, FP: False Positive, 0.05 is used as the cutoff. **B**, Receiver-operator characteristic (ROC) curves show HiNT performs better after the background subtraction. Areas under the ROC curves (AUCs) are shown in parentheses. **C**, The original, background (average of multiple other Hi-C maps), and the adjusted maps are shown for the three cases highlighted in panel A. Validated translocations are marked by circles.

Although the rank product approach detects the majority of translocated chromosomal pairs, four validated translocations are not identified. To investigate this issue, we compare the Hi-C interaction matrices of the detected (Supp. Fig. 8) and missed chromosomal pairs (Supp. Fig. 9). Compared to the detected chromosomal pairs, no translocation signature can be visually detected from the interaction matrices for missed pairs. In addition, the sharp boundaries at translocation breakpoints on the 1D coverage profile can only be found in our predicted translocated chromosomal pairs. Thus, we believe that there are some translocated chromosomal pairs that are simply not reflected within Hi-C data, or the validation data may be incorrect, e.g., due to the variation among the K562 lines. We further examined four more cancer cell lines, including HelaS3 (cervical carcinoma), LNCaP (prostate carcinoma), Panc1 (pancreatic carcinoma), and T47D (breast cancer). We found that the rank product and the maximum interaction perform better than the Gini index in LNCaP, T47D, and Panc1, whereas the rank product and Gini index are more predictive in HelaS3 (Supp. Fig. 7E).

### HiNT detects translocation breakpoints at single base-pair resolution using Hi-C chimeric reads

Once a chromosomal pair containing a translocation is identified based on the rank product, HiNT searches for the translocation breakpoint. For a translocation, the 1D row/column-sum profile should change abruptly at the breakpoint (Supp. Fig. 8, and Supp. Fig. 10A). To identify this point, we use a change point detection method called *breakpoints* from the R package *strucchange* [26], which adopts a linear model to detect one or several change points in multivariate time series. However, the majority of the change points detected by *breakpoints* are the result of lower mappability and unremoved compartment effects, and thus should not be identified as the translocation breakpoints (Supp. Fig. 10A). To remove these false positives, we impose a filtering step in which only those with one quadrant (unbalanced translocation) or two diagonally-opposed quadrants (balanced translocation) around the candidate breakpoint have very high interactions (Supp. Fig. 10, Methods). Here, we define a high interaction frequency as being greater than the 99th percentile of all the interactions between the two chromosomes.

Next, we determine the precise coordinates of the breakpoints by using *ambiguous* chimeric reads [16] (Fig. 1A). These reads have their primary alignment near a breakpoint in one chromosome (e.g. chrA) and their clipped part align near a breakpoint in another chromosome (e.g. chrB). HiNT provides the intervals in which the breakpoints occur (100kb resolution) and, as long as the breakpoint does not occur in an unmappable region, the exact location of the breakpoint (1bp resolution).

### Hi-C can supplement WGS by locating translocation breakpoints in repetitive regions

To assess its performance, we compare the translocation breakpoints determined from Hi-C using HiNT with those detected from WGS using Delly [27] and Meerkat [28]. In K562, 56 and 173 inter-chromosomal translocations are detected by Meerkat and Delly, respectively, with only 14 translocations detected by both (Fig. 5A). This level of discrepancy is not unexpected [29] and is indicative of the difficulty of detecting SVs in general. When we intersect these 14 consensus WGS-based translocations with those detected by HiNT, we find that 5 are in common (Fig. 5A). Two additional ones were found by HiNT and either Meerkat or Delly but not both. In these 7 cases, the breakpoints were exactly the same at the nucleotide level, confirming the accuracy of the calls (Supp. Table 2). An example is a translocation between chromosome 9 and 22 shown in Fig. 5B, with more than 100 supporting clipped reads in Hi-C data and many discordant reads in WGS data (Fig. 5C).

**Figure 5.**
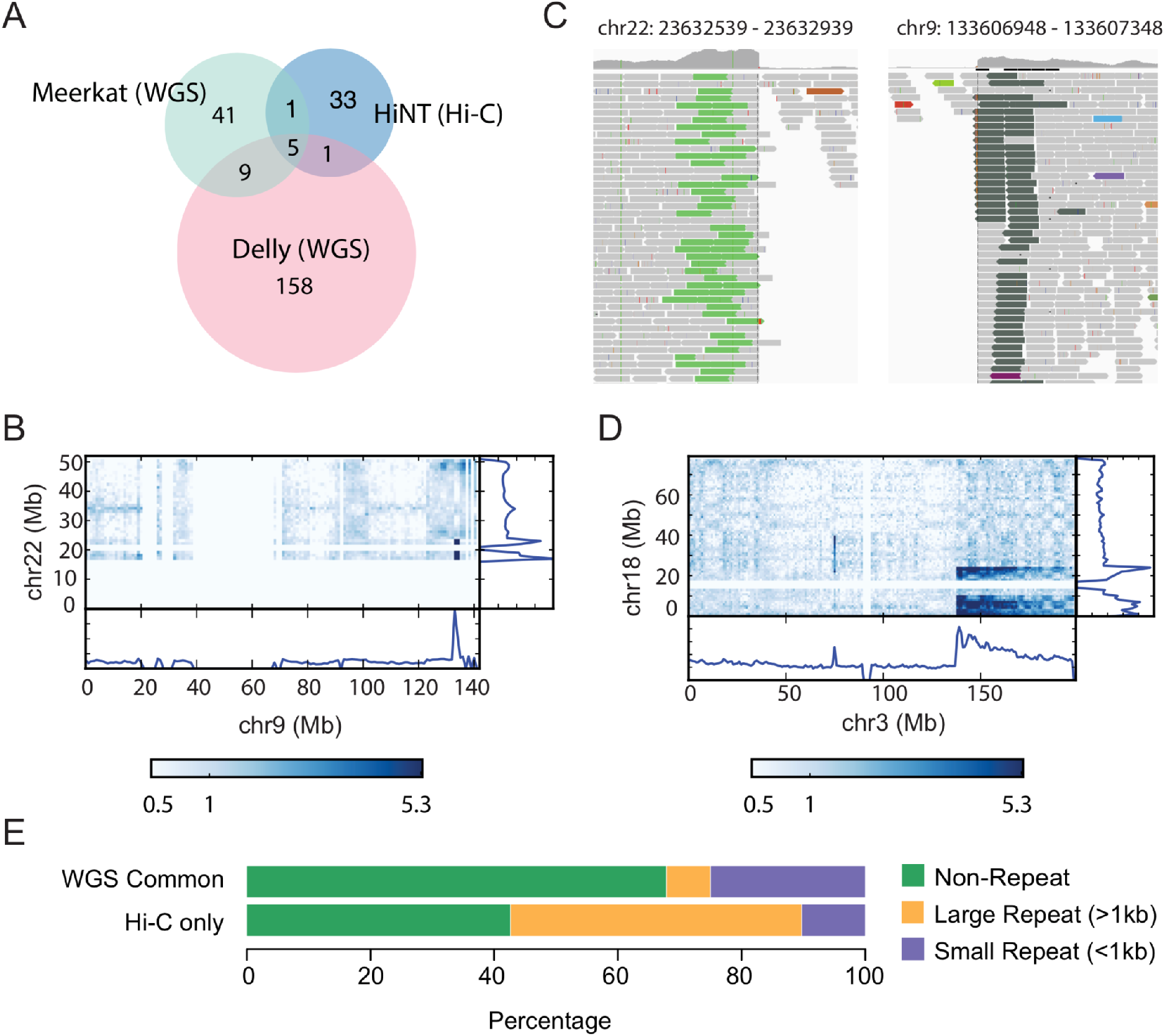
Comparison of breakpoints detected from Hi-C and WGS. **A**, Overlap of the translocation breakpoints detected by Meerkat (WGS), Delly (WGS), and HiNT (Hi-C). **B**, The Hi-C interaction map containing a breakpoint detected in both Hi-C and WGS. **C**, The same exact breakpoint in panel B is captured in WGS. Discordant reads in light green (dark green) are paired-end reads whose mates are found on chr9 (chr22). **D**, Hi-C interaction map illustrating a clear case of translocation detected only by HiNT. **E**, Breakpoints detected in both Meerkat and Delly (‘WGS Common’) and only in Hi-C only are classified into small repeat, large repeat and non-repeat regions, showing that Hi-C is enriched for SVs involving large repeats.

Thirty-three translocations are detected only from Hi-C data (Fig. 5A; listed in Supplementary Table 3 and can be viewed interactively by HiGlass [30] at http://18.215.251.253/). For example, a significant rank product score is found between chr3 and chr18 in the Hi-C interaction matrix (Fig. 5D), and three breakpoint regions are detected by HiNT including one validated by FISH [24] (Supp. Table 4). However, few discordant reads are identified from WGS. A major reason for this difference is the low mappability around those breakpoints. As illustrated in Supplementary Figure 1, the long physical distance between Hi-C read pairs allow identification of translocations whose breakpoints occur in a repetitive region—the paired reads can “jump over” the repeat region and map to surrounding mappable regions, even though the breakpoint itself cannot be mapped. Indeed, we find that large repeat (>1kb) regions (as found in repBase [31]) make up a disproportionately large fraction of regions containing Hi-C-only breakpoints compared to WGS consensus breakpoints (Fig. 5E). We note that repetitive regions with high sequence divergence are mappable, but we used the term ‘repetitive region’ for conceptual clarity.

For the translocations detected only in WGS, some are missed in Hi-C due to its more uneven genome coverage. In other cases, we find that, surprisingly, the discordant reads from WGS contain a large fraction of single nucleotide polymorphisms or have low mapping qualities, indicating issues in read alignment (Supp. Fig. 11). Consistent with that observation, translocation signatures are not found in the Hi-C interaction maps. These analyses suggest that Hi-C is a powerful tool to detect translocations and can complement WGS, especially for detecting those with breakpoints in repetitive regions.

### HiNT outperforms existing tools on detecting translocations

Others have attempted to identify structural variants from Hi-C data. One approach is simply to visually inspect the interaction heatmaps—a low resolution detection of breakpoints with poor scalability and reproducibility [23]. Better approaches search for regions that contain abnormal interaction frequencies based on normalized Hi-C interaction maps [6, 32]. However, such methods utilizing only contact frequencies cannot easily distinguish translocations from chromatin interactions, thus giving a high false discovery rate (FDR). A recent algorithm HiCtrans [19] identifies translocation breakpoints based on change-point statistics obtained by scanning the inter-chromosomal contact maps of each chromosomal pair. However, searching the breakpoints across all inter-chromosomal contact maps leads to a high computational cost.

For a comprehensive set of inter- and intra-chromosomal translocations, one could integrate WGS, Hi-C and optical mapping data [24]. However, in most cases, it is impractical to generate all these data types for a given sample. The method they used for Hi-C data [24] is hic_breakfinder, an iterative approach to locate local clusters that deviate from the expected interaction frequencies in a Hi-C contact matrix.

To compare the performance of these algorithms, we first apply HiCtrans [19] and HiNT to simulation data. Hic_breakfinder [24] is not used here because it requires the aligned reads in BAM format, but our simulation is matrix-based. Of the 21 simulated inter-chromosomal translocations (mix of balanced/unbalanced and heterozygous/homozygous translocations), HiNT identified 20 correctly while calling additional 5 breakpoints (Supp. Fig. 12A). The one missing translocation was located at the centromere of chr21 (Supp. Fig. 12B). In contrast, HiCtrans called 531 translocations (distributed across 100 different chromosomal pairs), but none were *bona fide* translocations (Supp. Fig. 12C).

We also compared HiNT, HiCtrans [19], and hic_breakfinder [24] on the K562 data. As shown in Supp. Fig. 12D-E, HiNT has the highest AUC measure (0.85 vs 0.78 and 0.77, see Methods) as well as the best precision-recall curve. Additionally, we found that while HiCtrans identified 132 translocated chromosomal pairs, which is more than half of the number of all chromosomal pairs, only 10 of them contain known translocations. Among all 931 breakpoints (~1Mb resolution) identified by HiCtrans, only 2 of them cover what are detected from WGS by both Meerkat and Delly (Supp. Fig. 12F). On the other hand, hic_breakfinder identified 77 breakpoints (~100kb resolution). Among these breakpoints, 4 are identified by both Meerkat and Delly (Supp. Fig. 12F). This suggests a higher false discovery rate of HiCtrans and hic_breakfinder than HiNT. Furthermore, we found that 60% (24/40) of HiNT-identified breakpoints can also be identified by other methods. In contrast, this value is only 35% (27/77) and 3.0% (28/931) for breakpoints output from hic_breakfinder and hictrans, respectively (Supp. Fig. 12F). Collectively, HiNT-TL outperforms HiCtrans and hic_breakfinder in both specificity and accuracy.

## Conclusion

Robust identification of SVs remains paramount to accurate inference of long-range interactions from Hi-C data. We have shown that HiNT can be used to identify CNVs and inter-chromosomal translocations with split read support for breakpoints whenever possible, and that it outperforms existing methods. Although not as sensitive as WGS data in general, Hi-C data can complement WGS data for detection of translocations in repetitive regions. As new technologies for capturing three-dimensional interactions are introduced, further computational methods will be needed to avoid the confounding effects of SVs.

## Methods

### Data sources

Hi-C data: *in-situ* Hi-C data in cancer cell line K562 and in normal cell lines including GM12878, HMEC, HUVEC, IMR90, and NHEK were obtained from GEO (Gene Expression Omnibus) with the accession number GSE63525 [16]. All the normal cell line data were combined to create the background Hi-C interaction matrix. Hi-C data for HelaS3, LNCaP, Panc1, and T47D, which were generated by the Dekker lab [33], were downloaded from the ENCODE website (See details from Supp. Table 1).

WGS data: We downloaded the BAM file for NA12878 WGS data from the 1000 genomes project [34], and the BAM file for K562 WGS data from the GDC legacy archive of the Cancer Cell Line Encyclopedia (CCLE) project [35].

### CNV identification from WGS

BIC-seq2 [36] was used to derive CNV segments from WGS read coverage data. For the segmentation step, we used *binsize* = 50,000 *bp* and *λ* = 50 to determine the final CNV breakpoints in NA12878. *λ* is a parameter that controls the smoothness (the number of breakpoints) of the final CNV profile. chrY and chrM were excluded from the analysis.

### Definition of copy ratios in Hi-C and WGS data

Copy ratio is defined as the ratio of observed and expected values. In Hi-C, observed values are the residuals from GAM Poisson regression, and expected values are set to zero. In WGS, observed values are read coverage, and expected values are estimated by a semi-parametric regression model via BIC-seq2 [36].

### Simulation of inter-chromosomal translocations in Hi-C contact maps

The simulation pipeline defines two random coordinates from distinct chromosomes as the origin and destination of the translocation (e.g. x on chr1, and y on chr2). Then, it creates the translocated version of interaction matrices for chr1 to chr1, chr2 to chr2, and chr1 to chr2 via rearranging the original interaction probabilities.

### SV detection from WGS

SV detection from WGS was carried out using Delly and Meerkat. To omit germline SVs, we used NA12878 as a control genome. Default parameters were used to run Delly. Only translocations that passed the internal quality control and were marked as “PRECISE” in Delly were used for comparison. Default parameters were used to run Meerkat, and filtering step was performed according to the post-processing steps described in the tool manual. Only valid precise inter-chromosomal translocations were kept for comparison.

### Gini Index calculation

For each Hi-C inter-chromosomal interaction matrix M, we first sorted the contact regions, based on the adjusted contact frequencies between these two regions, from lowest to highest, then calculated the cumulated contact frequencies of matrix M. Regions that did not form contacts with any other regions were excluded. A plot of this functional relationship is called a Lorenz curve. The Gini index is computed as twice the area between the Lorenz curve and the diagonal.

### Breakpoint filtering

To remove false discovered change points, we first construct two-dimensional Cartesian coordinate systems originating from the intersection of each pair of candidate breakpoints. For each coordinate system, we then define four, 5-bin-by-5-bin quadrants around the origin, and we calculate the average interaction frequency within each quadrant (Supp. Fig. 10A). The valid breakpoints for translocations should have only one (unbalanced translocation) or two (balanced translocation) quadrants with very high interactions, and the remaining quadrants should have lower interaction frequencies (Supp. Fig. 10B upper panel). More specifically, for balanced translocations, the two quadrants with high interaction frequencies should diagonally oppose each other (Supp. Fig. 10B upper panel). If zero, three, or all quadrants have high interaction frequencies, the proposed breakpoints are considered false positives and removed (Supp. Fig. 10B lower panel). Here, we define a high interaction frequency as being greater than the 99th percentile of all the interactions between the two chromosomes.

### ROC curves of HiCtrans and HiC_breakfinder on translocated chromosomal pairs prediction

To create ROC curves for the evaluation of translocated chromosomal pairs prediction, we rank all the chromosomal pairs first. Both HiCtrans and hic_breakfinder output a score (entropy ratio in HiCtrans, and log-odds in hic_breakfinder) to measure the strength of each breakpoint call.

We assign each chromosomal pair a representative score by taking the score of the most significant breakpoint that located in this chromosomal pair. The chromosomal pairs are then ranked by the representative scores. ROC curves and AUC values are calculated by using the R package *ROCR* [37]. The chromosomal pairs reported in the literature or validated by FISH experiments are used as true positives here.

### Details of the HiNT pipeline

1. HiNT-PRE: Raw Hi-C data in FASTQ format are aligned to a reference genome (hg19) via bwa-mem: `bwa-0.7.16a-r1185-dirty/bwa mem-SP5M bwaIndex/hg19.fa inl.fq in2.fq`. Read pairs that are both uniquely mapped to the genome are collected as valid pairs. However, 10%-20% of the remaining Hi-C read pairs contain at least one chimeric read with split alignments. Chimeric pairs with one read uniquely mapped and the other chimeric, due to ligation, are defined as *unambiguous* chimeras [16], and counted as valid pairs. All other chimeric pairs are classified as *ambiguous* [16] chimeras, and are used to identify translocation breakpoints at single base-pair resolution. All the unmapped, multi-mapped, and PCR duplicated read pairs are discarded from our analysis. All pairs are classified by pairtools (https://github.com/mirnylab/pairtools). Then, a Hi-C interaction matrix is generated from all the valid pairs by cooler [22] or juicer tools [38] at 50kb, 100kb, 1Mb, or at a user-specified resolution.

2. HiNT-CNV: First, a 1D coverage profile for each 50kb bin (default) is calculated along the whole genome using an unnormalized contact matrix. Bin size can be specified by users based on the sequencing depth and accuracy need. Then, a GAM regression with a Poisson link function is performed to remove the known Hi-C biases with pre-calculated GC content, mappability, and the number of restriction sites in each bin. Then, the segmentation method of BIC-seq is applied to the regression residuals to identify the breakpoints and generate the final CNV profile.

3. HiNT-TL: Translocation detection is performed in three steps; determination of the translocated chromosomal pairs, identification of the rough breakpoint regions, and determination of the exact breakpoints at single base pair resolution. To determine the translocated chromosomal pairs, 1 Mb-binned and genome-wide normalized inter-chromosomal interaction matrices are taken as input. To remove the effects of A/B compartments, a background model is created by averaging multiple *in-situ* Hi-C data in normal cell lines (Supp. Table 1). Each inter-chromosomal interaction matrix is corrected with the background by taking the ratio between the original signals and the background signals. Then, for each possible chromosomal pair, Gini index and the maximum contact frequency are calculated. Then, a rank product score is computed *RP_i_* = (*R_gini,i_/n*) * (*R_mif,i_/n*), where *R_gini,i_* and *R_mif,i_* are the ranks of matrix *i* based on Gini index and maximum interaction frequency, respectively, and *n* is the total number of inter-chromosomal interaction matrices. Chromosomal pairs with *RP_i_* ≤ 0.05 are defined as the potential translocated chromosomal pairs.

HiNT then calculates the 1D coverage profiles by calculating the sum of each row and column of the adjusted inter-chromosomal interaction matrices for those predicted translocated chromosomal pairs. It then applies the function *breakpoint* in the R package *strucchange,* a function with high computing performance that allows simultaneous estimation of multiple breakpoints in a given time series data, to the coverage profiles to identify all change points. The translocation rough breakpoint regions are further decided after the filtering step as we described in Supp. Fig. 10.

To get the precise breakpoints at single base-pair resolution, HiNT uses the soft-clipped reads-based algorithm that is commonly used for WGS SV prediction. Translocation breakpoints that are covered by at least one split read pair with one end mapped to the rough breakpoint region on one chromosome, and the other end mapped to the rough breakpoint region on another chromosome are reported at single base-pair resolution; otherwise, the predicted rough breakpoint regions will be reported. Not all the breakpoints are expected to have supported clipped reads due to the non-uniform distribution of read coverage in Hi-C data.

## List of abbreviations

HiNT: **Hi**-C for copy **N**umber variation and **T**ranslocation detection;
CNV: copy number variation;
SV: structural variation;
GAM: generalized additive model;
WGS: whole genome sequencing;
1D: 1-dimensional;
ROC: receiver operating characteristic;
TADs: topologically associated domains;
TP: true positive;
TN: true negative;
FP: false positive;
FN: false negative;
RP: rank product.

## Declarations

### Availability of data and material

HiNT is available open source at https://github.com/parklab/HiNT.

Sources of the data used in this study are included in Supp. Table 1

### Authors’ contribution

P.J.P., B.A., and S.W. conceived the project and method design. S.W. processed the data and implemented HiNT. S.L., C.C., D.J., J.W., G.N., B.A., and P.J.P. discussed and helped to implement HiNT. S.W. and P.J.P. wrote the manuscript with assistance from the other authors.

All authors read and approved the final manuscript.

## Acknowledgements

We thank Shannon Ehmsen for preparing the figure of HiNT workflow.

