## Supplementary Figures for "HiNT: a computational method for detecting copy number variations and translocations from Hi-C data"

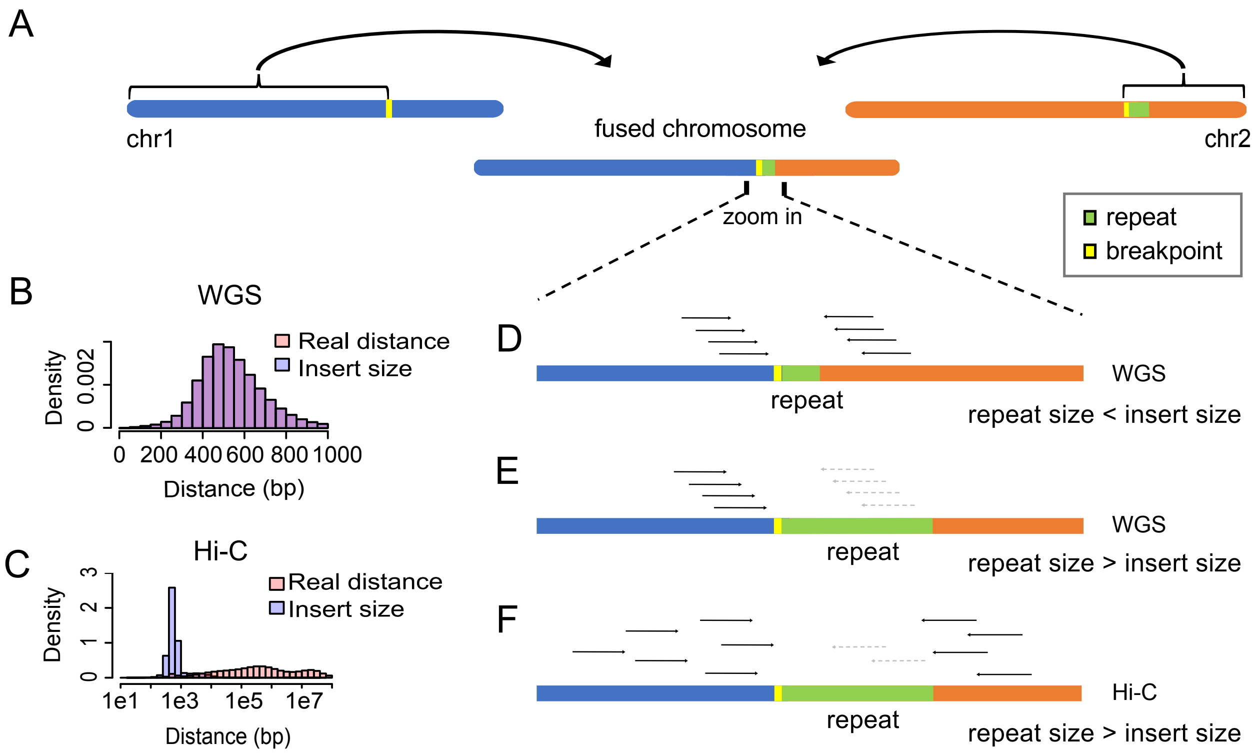


**Supplementary Figure 1.** Hi-C data is superior to WGS in variation detection in repetitive regions. **A,** Illustration of a fused chromosome with a breakpoint located in repetitive region. **B-C,** The distribution of the real distances (pink) between two mates, and the insert sizes (light blue) in WGS (B) and Hi-C (C). **D,** Reads can be correctly mapped to the reference genome if repeat size is less than the insert size in WGS. **E,** Reads cannot be correctly mapped to the reference genome if repeat size is larger than the insert size in WGS. **F,** Reads surrounding the repetitive regions can be used to detect the breakpoint in Hi-C.

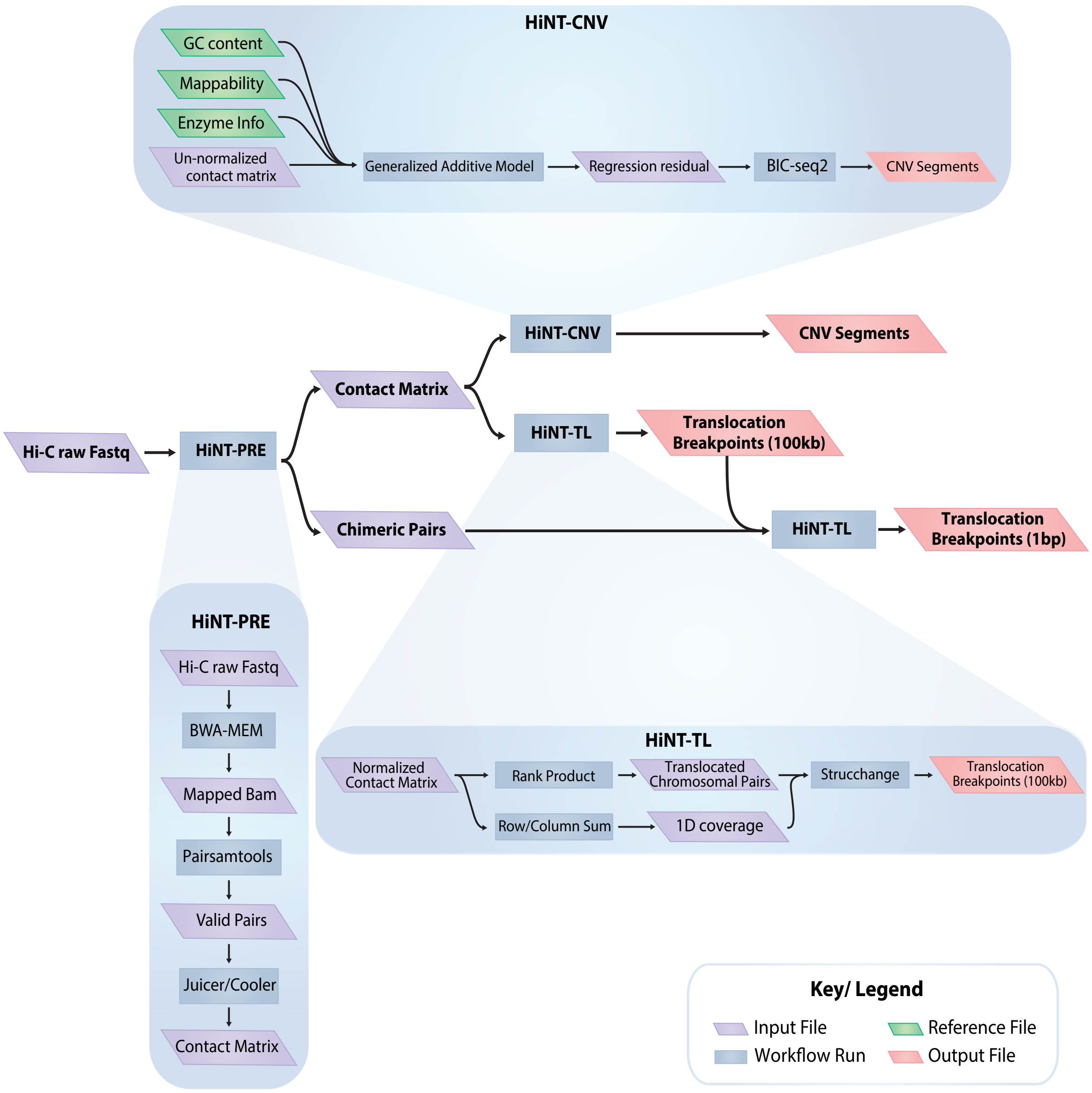


**Supplementary Figure 2.** Overview of the HiNT workflow. HiNT has three components: HiNT-PRE, HiNT-CNV, and HiNT-TL. HiNT-PRE preprocesses Hi-C data to generate the contact matrix; HiNT-CNV performs CNV detection; and HiNT-TL detects translocation breakpoints at 100kb as well as base-pair resolution.

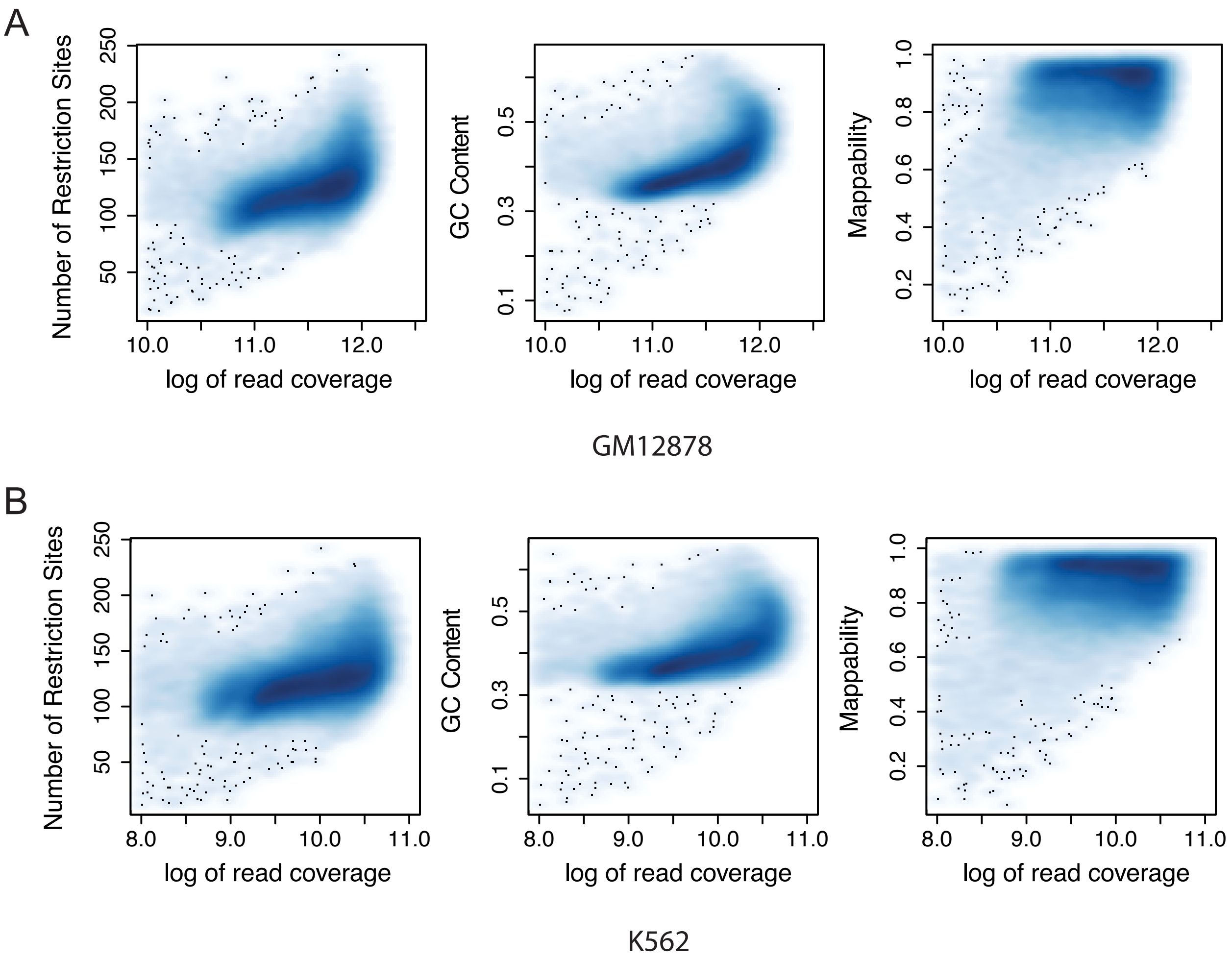


**Supplementary figure 3**. Correlation between the natural log of 1D coverage and the number of restriction sites (left), GC content (middle), and mappability (right) in each 50kb bin in GM12878 (A) and K562 (B) cell.

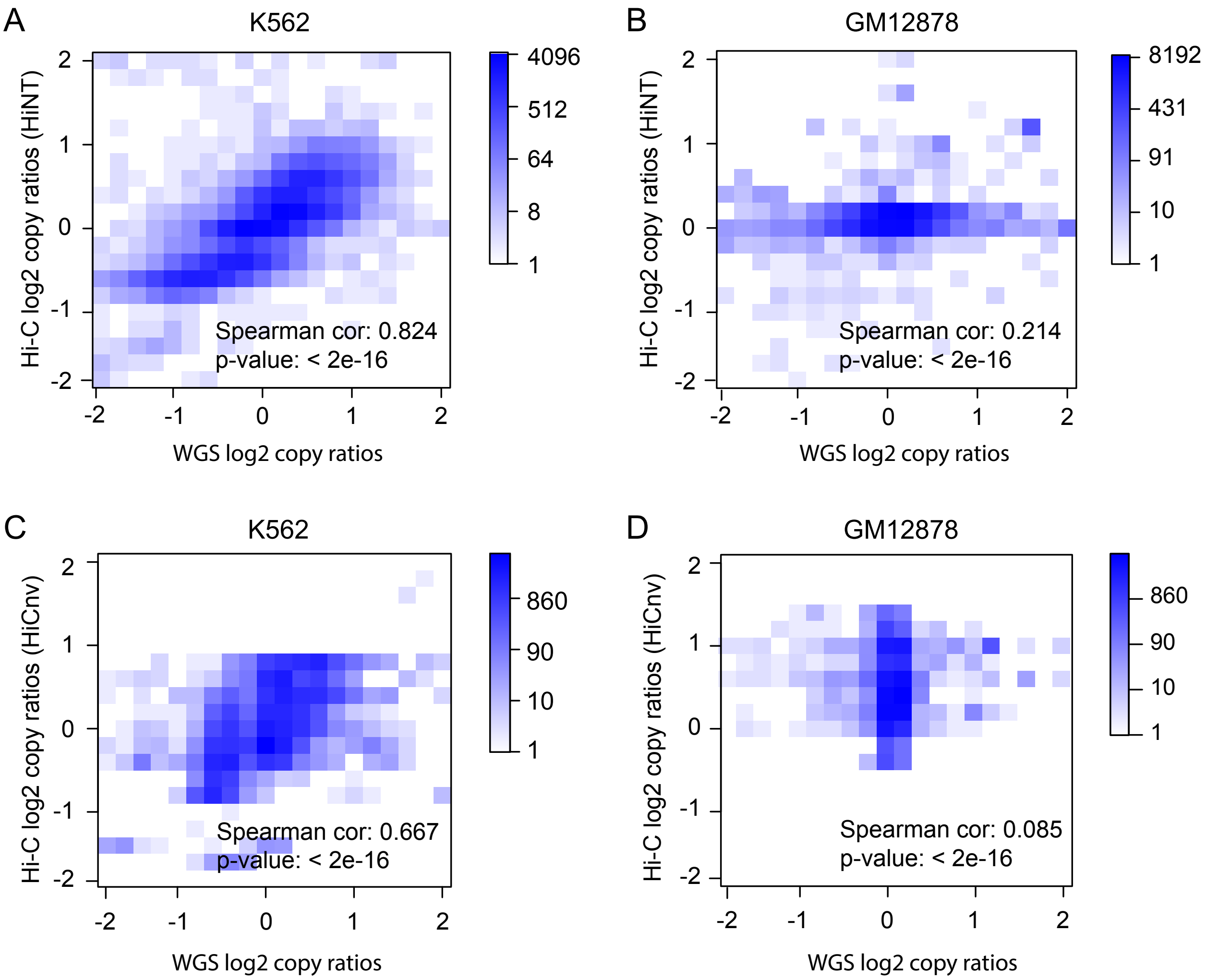


**Supplementary figure 4**. CNVs detected by HiNT from Hi-C are consistent with those detected from WGS. **A-B,** Correlation of log2 copy ratios in each bin (50kb) detected from WGS and Hi-C (HiNT) in K562 (A) and GM12878 (B). **C-D,** Correlation of log2 copy ratios in each bin (50kb) detected from WGS and Hi-C (HiCnv) in K562 (C) and GM12878 (D).

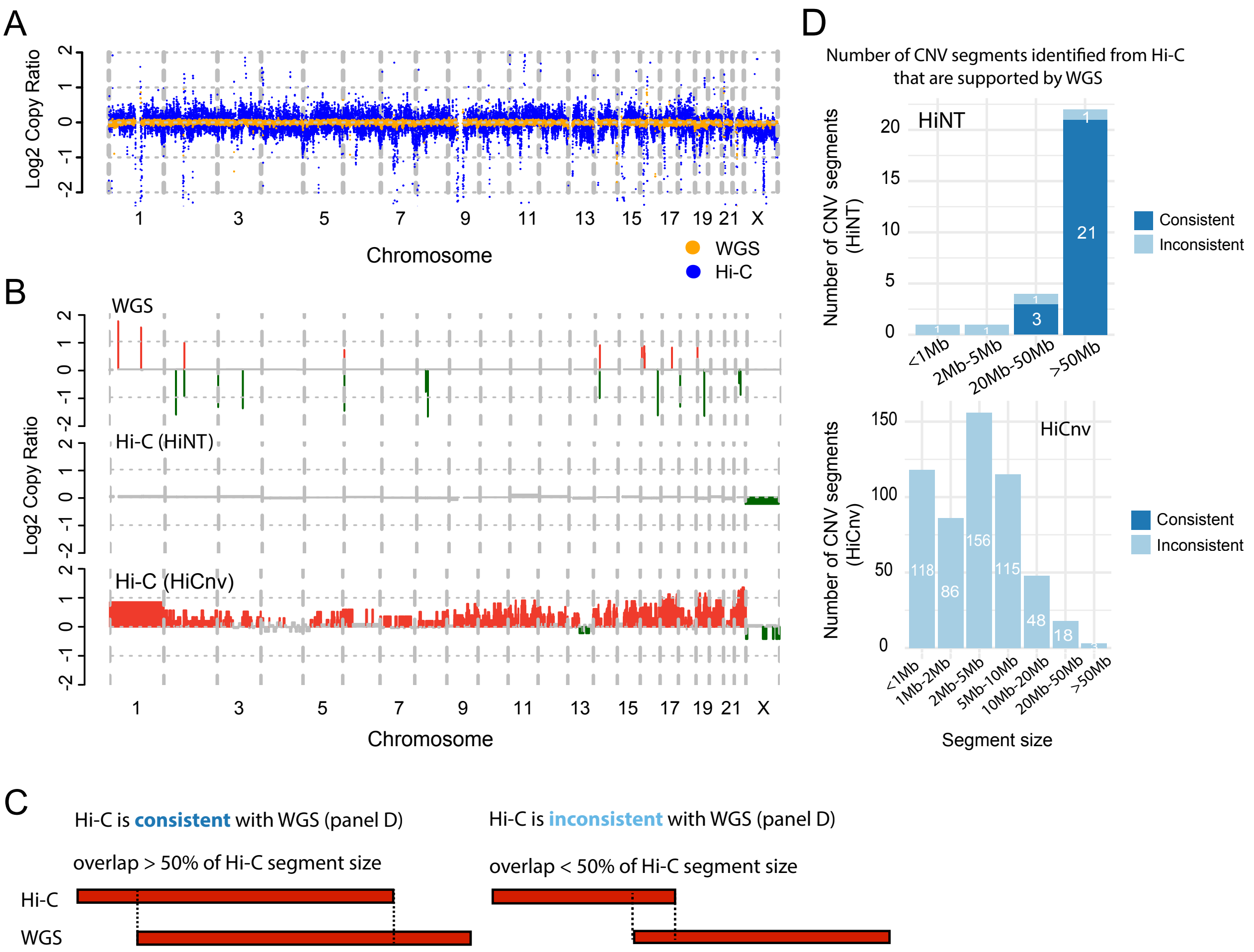


**Supplementary figure 5**. Copy number inference in GM12878 cells by HiNT. **A,** Comparison of log2 copy ratios calculated using regression residuals from Hi-C (blue) and using read coverage from WGS (orange). **B,** Comparison of CNV profiles from Hi-C and WGS after segmentation. Red, green and grey bars represent copy gain, copy loss, and copy neutral regions, respectively. **C,** Schematic of the consistency analysis. CNV segment detected from Hi-C is consistent with that detected from WGS if the overlapped region is larger than 50% of the original segment size, and vice versa. **D,** The number of CNV segments detected from Hi-C by HiNT (upper) and HiCnv (lower) that are also supported by WGS.

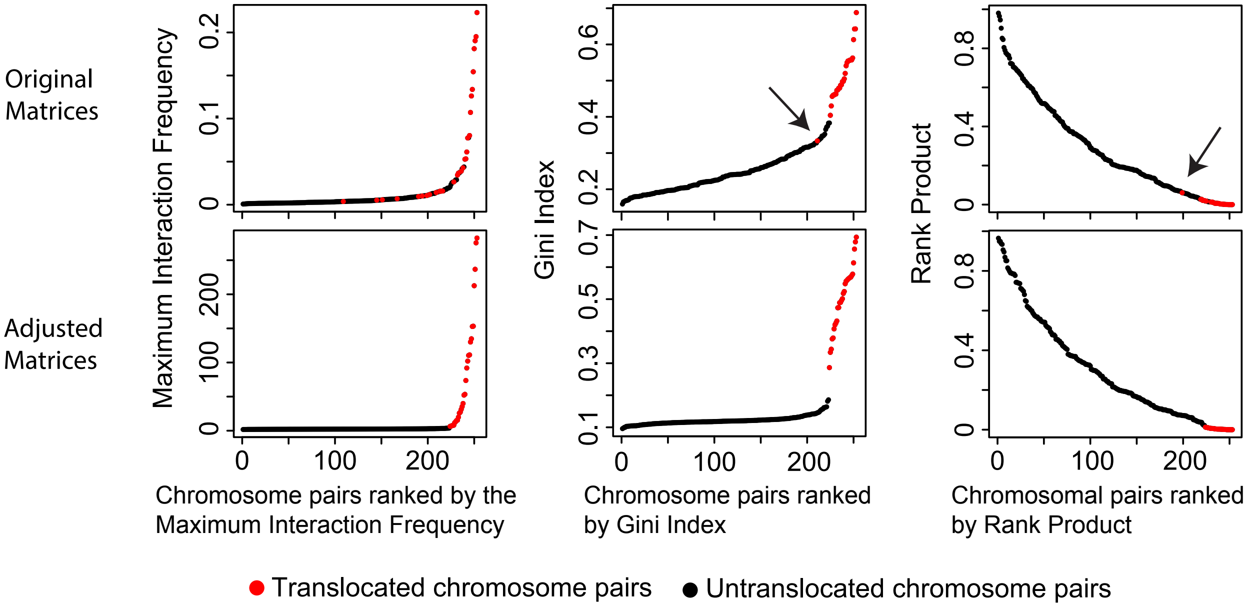


**Supplementary Figure 6.** Rank Product approach accurately identifies simulated translocated chromosome pairs. Distribution of the maximum interaction frequency (left), the Gini Index in an inter-chromosome contact matrix (middle), and the rank product of these two (right) in Hi-C data with simulated translocations.

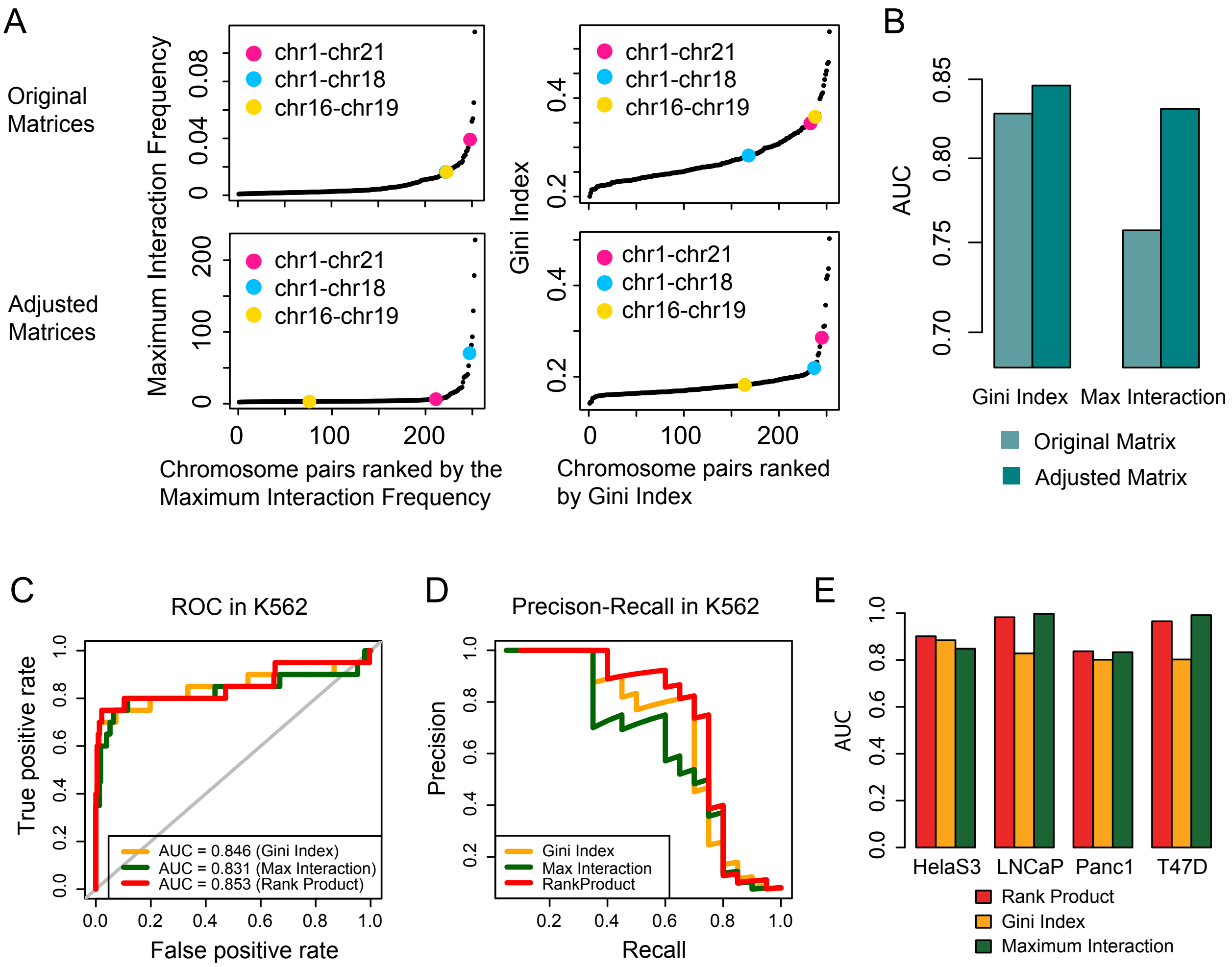


**Supplementary Figure 7.** Rank Product approach accurately identifies translocated chromosome pairs in K562 cells. **A,** The distribution of the maximum interaction frequency (MIF, left), the Gini Index (right), and the rank product of these two (figure 4A) in inter-chromosome contact matrices before (upper) and after (lower) adjustment in K652. Chromosomal pairs in pink and blue correspond to two FISH-validated translocation pairs (chr1, chr21) and (chr1, chr18); the one in yellow corresponds to a chromosome pair (chr16, chr19) without translocation. **B,** AUROC values show either Gini Index or MIF perform better after the background subtraction. **C-D,** ROC curves (C) and precision-recall curves (D) of translocated chromosomal pairs predicted by using Gini Index only (orange), the maximum interaction only (dark green), and the rank product of these two (red) in K562. **E,** Performance of rank product, Gini index, and the maximum interactions in HelaS3, LNCaP, Panc1, and T47D.

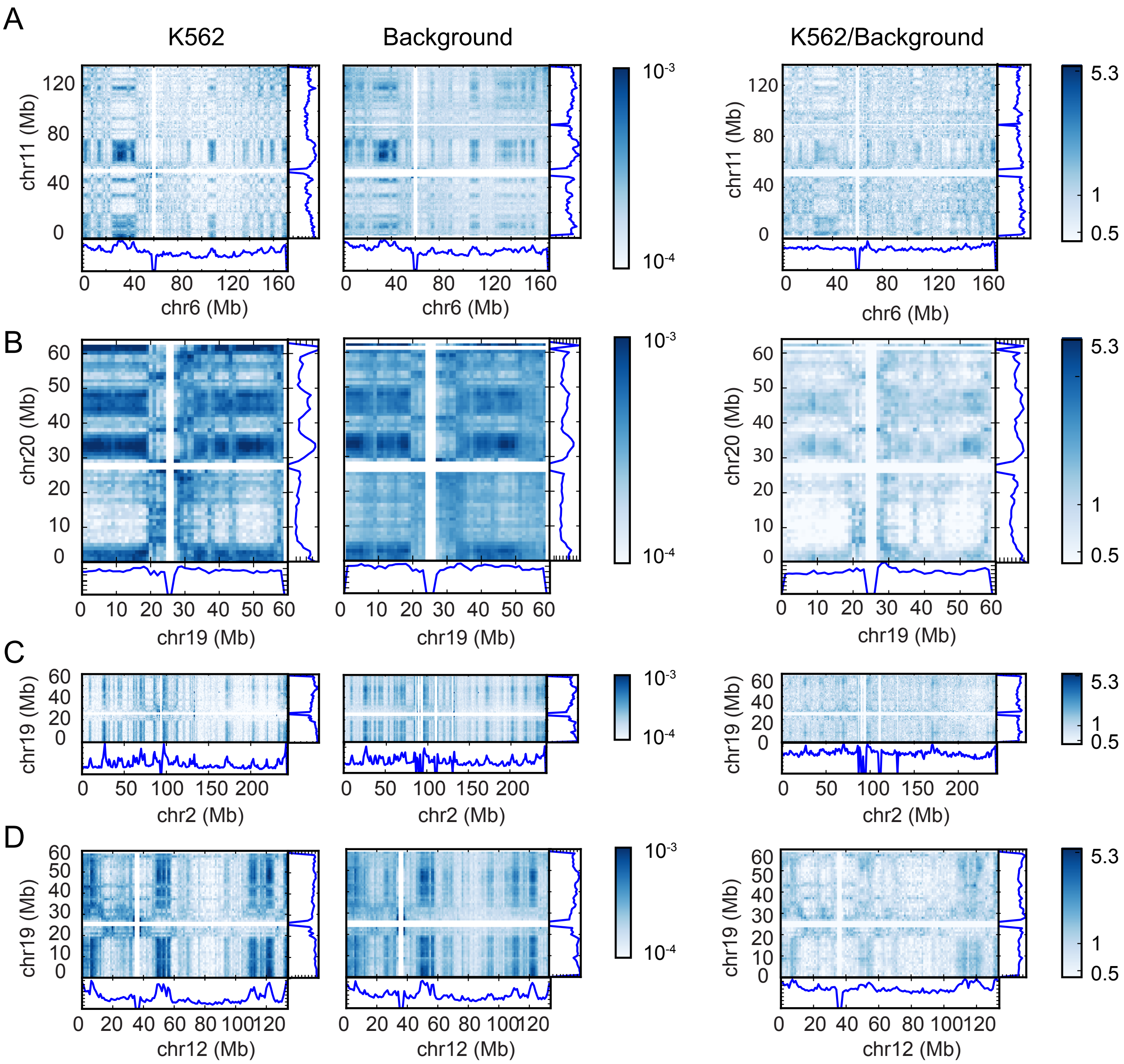


**Supplementary Figure 8**. Examples of chromosomal pairs with most significant rank product. **A-D**, Hi-C inter-chromosomal heatmaps and 1D coverage in original K562(left), background (middle), and adjusted K562 (K562/Background, right) data.

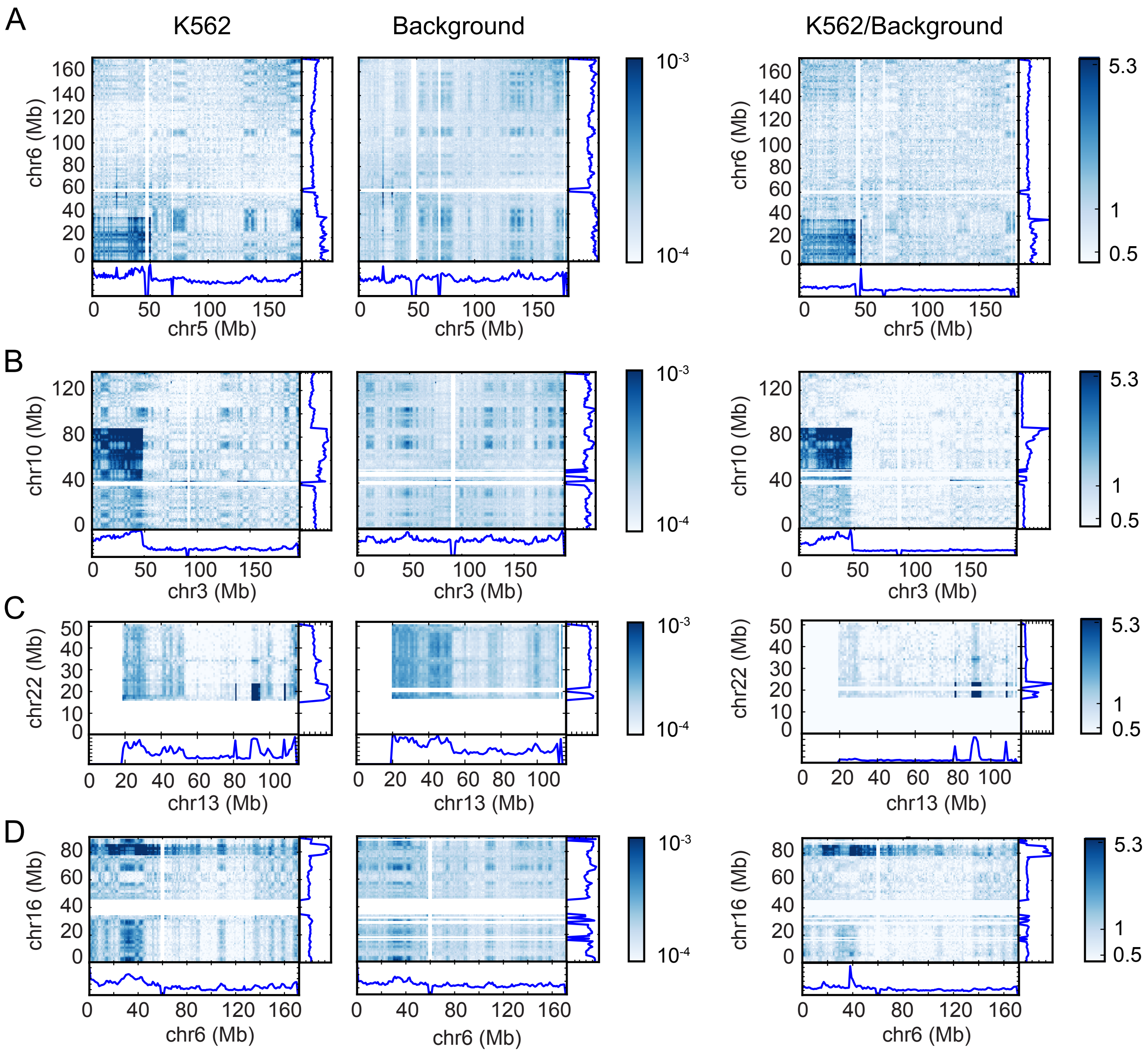


**Supplementary Figure 9**. Examples of missed translocated chromosomal pairs by HiNT. **A-D**, Hi-C inter-chromosomal heatmaps and 1D coverage in original K562(left), background (middle), and adjusted K562 (K562/Background, right) data.

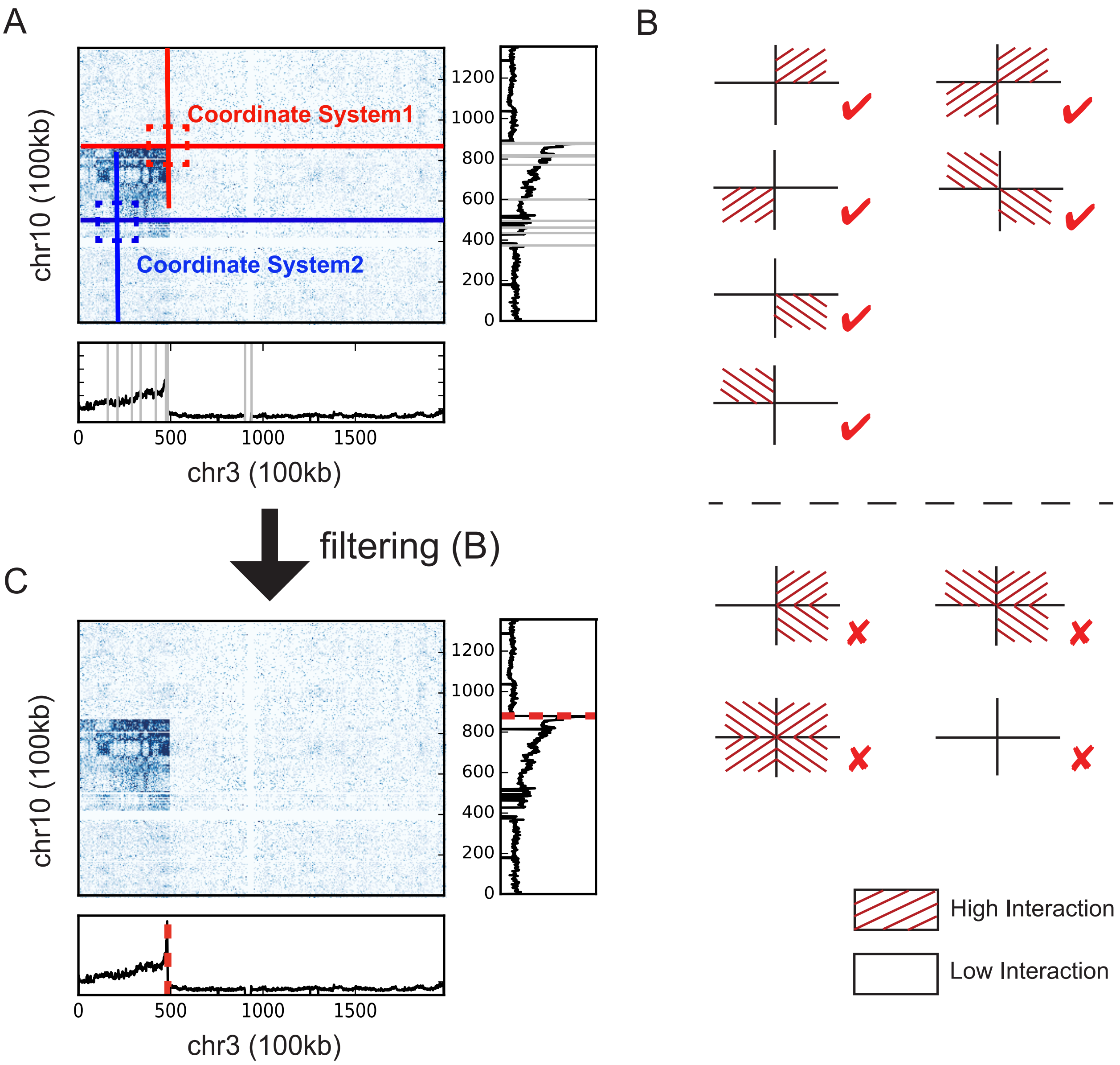


**Supplementary Figure 10**. Breakpoint detection and filtering. **A,** Candidate breakpoints (grey lines) detected by *strucchange* based on the 1D coverage profile. Two-dimensional Cartesian coordinate systems originating from the intersection of each pair of candidate breakpoints are constructed; two examples are shown in the figure. **B,** Patterns of Hi-C interaction frequencies in four 5-bin-by-5-bin quadrants, that generated by the pair of breakpoints from both chromosomes. Valid translocation breakpoints are shown above the dash line, and invalid breakpoints are shown below. **C,** Translocation breakpoints (red dotted lines) after the filtering step.

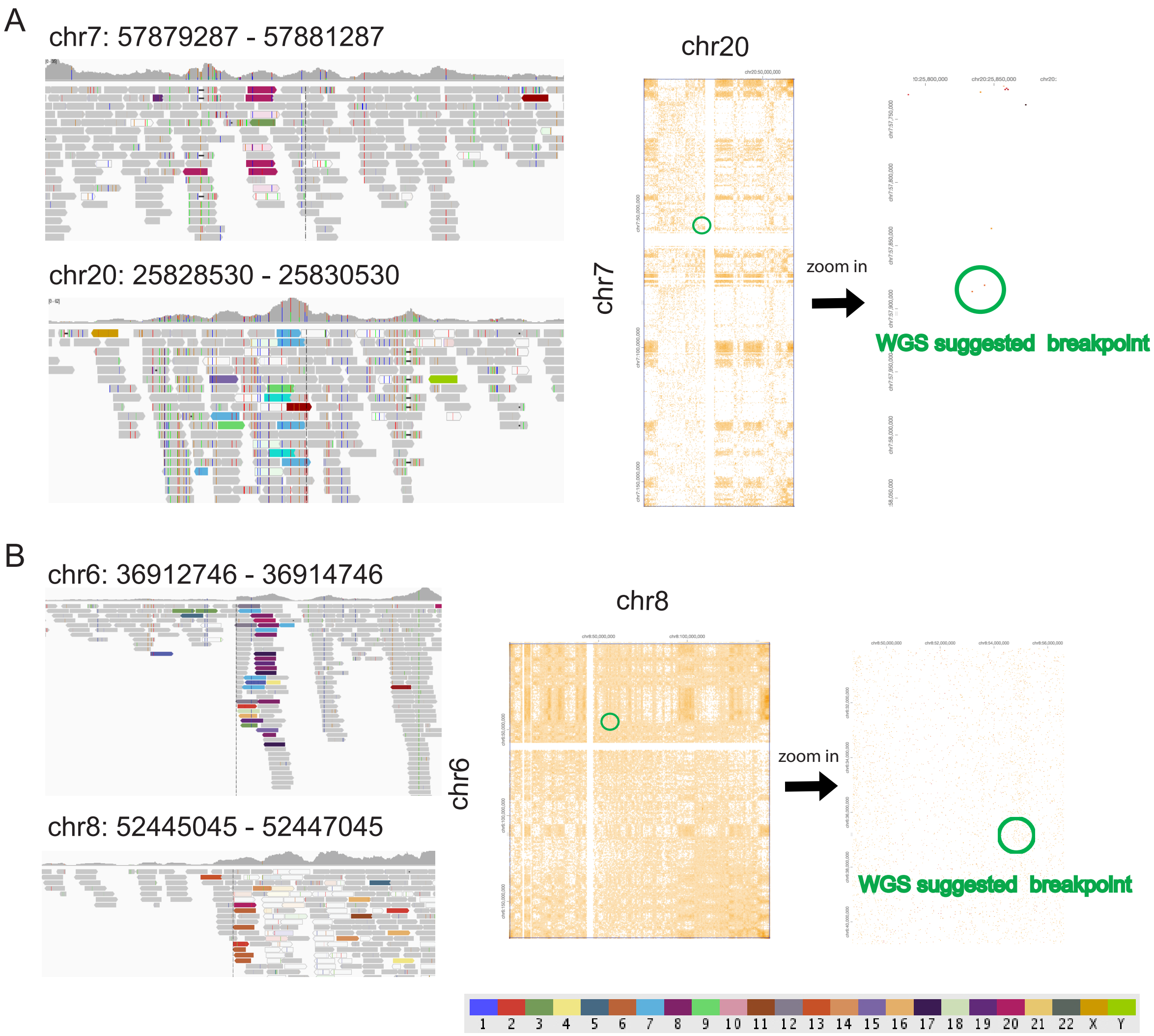


**Supplementary Figure 11.** Examples of the translocations detected from only WGS. **A,** The distribution of discordant reads and clipped reads around the translocation breakpoints detected from WGS on chr7 and chr20 (left); Hi-C interaction heatmap across the whole chromosomes (middle) and regions around breakpoints (right). **B,** Similar to A, but the translocation between chr6 and chr8. In the IGV screenshot, each color bar represents a SNV (single nucleotide variant), and the colored reads are paired end reads coded by the chromosome on which their mates can be found. The color code is shown at the bottom.

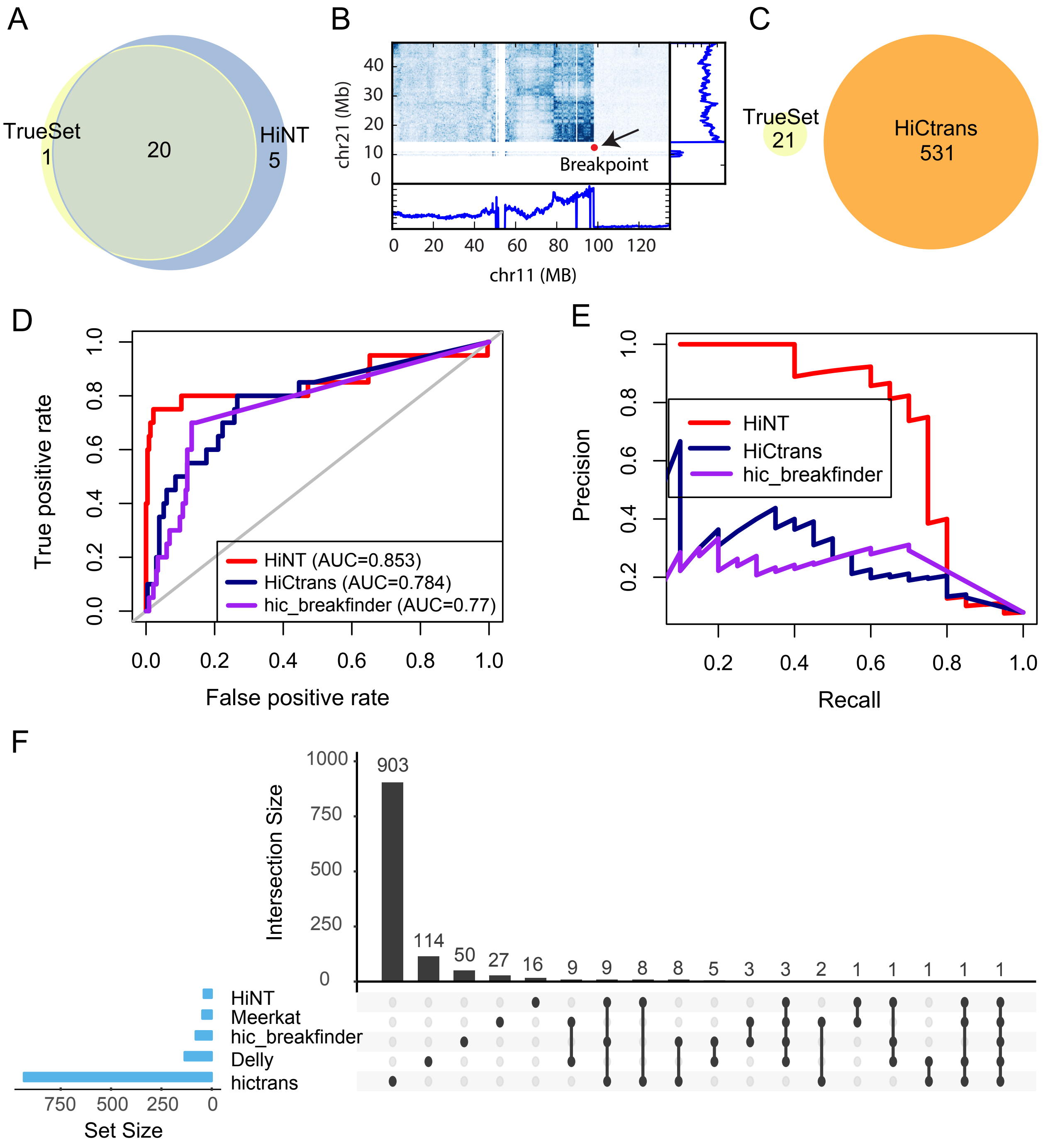


**Supplementary Figure 12**. HiNT outperforms existing methods on translocation breakpoints detection in both simulated and real Hi-C data. **A,** The overlap of translocation breakpoints detected by HiNT and simulated true set. **B,** Hi-C interaction heatmap for the breakpoint that was missed by HiNT. **C,** The overlap of translocation breakpoints detected by HiCtrans and simulated true set. **D-E,** Evaluation of the performance of HiNT, HiCtrans, and hic_breakfinder on translocated chromosome pairs prediction in K562 by ROC curves (D) and precision-recall curves (E). **F,** Intersections of the translocation breakpoints detected by Meerkat and Delly from WGS, and HiNT, HiCtrans and hic_breakfinder from Hi-C.
